# EEG entropy reflects both intrinsic and stimulation-induced corticospinal excitability

**DOI:** 10.1101/2025.05.16.654439

**Authors:** Simon Ruch, Joel Frohlich, Marius Keute, Ge Tang, Nelli Keksel, Alireza Gharabaghi

## Abstract

**Background:** Cortical excitability fluctuates throughout the day on multiple timescales, ranging from milliseconds to hours. This is reflected in the large variability of the brain’s response to transcranial magnetic stimulation (TMS). However, robust and interpretable biomarkers of the brain’s current excitability state are lacking.

**Objective:** We investigated whether entropy derived from singular value decomposition of short electroencephalography (EEG) segments could serve as a biomarker of cortical excitability as probed by TMS.

**Methods:** Entropy was computed from 1-second EEG segments preceding single-pulse TMS applied over the motor cortex. We assessed whether spontaneous fluctuations in pre-pulse entropy predicted trial-by-trial variability in TMS-induced motor-evoked potentials (MEPs). Additionally, we evaluated whether entropy tracked stimulation-induced changes in cortical excitability.

**Results:** Higher pre-pulse entropy, particularly over frontal regions, was associated with larger MEP amplitudes. TMS locally increased entropy over the motor cortex, while entropy decreased in other regions during the intervention. Participants who showed greater local increases in entropy from pre-to post-intervention also demonstrated larger increases in corticospinal excitability.

**Conclusion:** Entropy derived from short EEG segments reflects both intrinsic and stimulation-induced changes in cortical excitability. This marker may help optimize TMS interventions by informing brain state-dependent stimulation strategies and providing an index of intervention efficacy.

## Introduction

Cortical excitability fluctuates throughout the day at both short and long timescales. For example, the magnitude of cortical excitability as measured with transcranial magnetic stimulation (TMS) is modulated at the single-trial level (short timescale) by spontaneous ongoing neuronal oscillations in the slow-wave (<1 Hz), alpha (8-12 Hz), and beta (15 – 25 Hz) range [1–5]. Beyond these short period fluctuations, cortical excitability is also modulated by circadian rhythms (intermediate timescale; [6,7]), metabolic state [8,9], and sleep-deprivation (long timescale; [10,11]).

Currently, there are no widely-agreed upon markers that quantify the current state of cortical excitability at a global or local level and that can map fluctuations in excitability at both short and long timescales [12]. Markers for real-time estimation of the brain’s current state of excitability might help improve interventions, for example by providing targets for brain state-dependent TMS in the treatment of motor symptoms after stroke [13,14]. Furthermore, such markers may allow for a direct assessment of the excitability changes induced by an intervention [15].

Here, we ask whether measures of EEG signal entropy provide markers for the brain’s state of excitability [16]. Measures of entropy quantify the degree of complexity – also referred to as irregularity, variability, or randomness [17] – of a signal. Although different entropy measures derive from different theoretical backgrounds such as information theory or thermodynamics [18,19], most of them share similar properties [20,21]. They capture both trait-and state-like aspects of the EEG signal [22], reflect the 1/f shape of the EEG power spectrum [23], and map the balance between excitatory and inhibitory brain activity [16]. Entropy measures have the benefit that they often map non-linear dynamics in the EEG without an explicit model [24], provide interpretable values that may not need baseline correction, and do not require predefining specific frequencies of interest [25].

In the present work, we used the singular value decomposition (SVD) entropy [26] computed from scalp EEG data for estimating the local state of cortical excitability. We chose this specific measure because in-house tests showed that it has relatively fast computation time and is robust even when calculated over short EEG segments. Previous studies have successfully used SVD entropy for detecting schizophrenia [27], epileptic seizures [28], Alzheimer’s disease [29] and Parkinson’s disease [30], measuring the depth of anesthesia [31] and sleep [32], and detecting perceptual awareness [33].

We investigated whether SVD entropy calculated for short, surface Laplacian transformed EEG time series can assess the current local state of cortical excitability. To this end, we tested a single-pulse TMS intervention applied over the motor cortex. Specifically, we investigated whether inter-trial fluctuations in pre-pulse entropy were associated with TMS-induced motor evoked potential (MEP) amplitudes at the single-trial level over approximately 1h. We took advantage of the fact that the MEP amplitudes vary from pulse to pulse as a function of spontaneous fluctuations in brain excitability from trial to trial [34]. We further investigated how TMS-induced changes in corticospinal excitability relate to induced changes in entropy.

## Methods

The data reported here were obtained in a study in which participants underwent eight sessions of single-pulse TMS targeting the left motor cortex. In each session, 300 single TMS pulses were applied at either 100% (low intensity) or 120% (high intensity) of the resting motor threshold (RMT). The MEP was recorded at the right extensor digitorum communis (EDC) muscle. Before and after the intervention, resting state EEG measurements were performed. Furthermore, corticospinal excitability (CSE) was measured separately at 100% and 120% RMT intensity (hereafter, CSE100/120) before (pre) the intervention, immediately after the intervention (post1), and after a delay of 30 minutes (post2).

### Participants

Twenty healthy adults (11 female) participated in this study. Mean age was 27 ± 4.4 years (median 26; range 19 to 37). All participants were right-handed, reported no regular medical or recreational drug intake at the time of participation, and had no history of neurological, neurosurgical, or psychiatric treatment. An assessment of the inclusion and exclusion criteria was carried out via questionnaire before the intervention. The study protocol conformed to the Declaration of Helsinki and was approved by the Ethics Committee of the Medical Faculty at the University of Tübingen. The study followed the current safety guidelines for application of TMS [35]. No participant reported side effects. All participants gave written informed consent prior to the experiment.

Three participants were post hoc excluded from analyses due to poor EEG data quality (N=2) or data loss (N=1) in multiple sessions. The final sample thus consisted of seventeen participants.

### General procedure

We conducted a double-blind, randomized within-subjects experimental study. Each subject took part in eight sessions of single-pulse TMS targeting the left motor cortex that were scheduled at least 48 hours apart and around the same time each day. In each session, participant received 300 TMS pulses over the left primary motor cortex with a jittered inter-stimulus-interval (ISI) between 3.5 and 4.5 s. The exact timing of TMS pulses was controlled by the oscillatory phase of an EEG signal that was measured and processed in real time over bipolar electrodes approximately 1 cm posterior and anterior to electrode C3. The primary readout was the muscle response at the right EDC muscle. Across sessions, we varied TMS intensities (100% RMT, 120% RMT), target frequencies (10 Hz, 20 Hz), and target phases (-90°, 90°) in a fully crossed design. Two additional sham sessions were applied which stimulated the right shoulder (100% RMT and 120% RMT). The data from sham sessions are not reported here. The impact of phase-targeting will be reported in another manuscript.

To keep participants alert, 120 trials of a psychomotor vigilance task (PVT) were pseudo-randomly interspersed between TMS pulses, with the same ISI distribution. Participants were instructed to keep their gaze on a grid of red LED lights placed ∼50 cm in front of them. In PVT trials, the LED grid flashed for ∼50 ms. Participants were instructed to press a key on a custom-made input device with their left index finger as fast as possible, and reaction time (measured in ms) was fed back visually.

At the beginning of each intervention session, participants were seated comfortably in a reclining chair. The EDC hotspot and RMT were determined following standard procedures (see below). Six minutes of resting data were collected, with alternating 30s epochs of eyes open and eyes closed. A number of assessments were carried out before (pre-assessment) the intervention, as well as immediately following the intervention (post1) and following an additional delay of 30 minutes (post2). The TMS interventions comprised 300 single TMS pulses at 100% or 120% RMT intensity that were targeted at the peak or trough phase of ongoing 10 Hz or 20 Hz oscillatory activity. Intensity, as well as target phase and frequency were varied between sessions, but within participants, yielding a 2×2×2 design. Note that for the research question addressed here, the conditions target phase (peak/trough) and target frequency (10 Hz / 20 Hz) were not analyzed. The four sessions within each intensity (100% and 120%) were treated as independent recordings within each participant. Participants were asked to keep their right hand completely relaxed, while passively sitting and fixing their eyes on a fixation cross displayed on a monitor. The pre-and post-intervention assessments comprised of four protocols aimed at assessing the impact of the interventions on changes in CSE measured separately at 100% and 120% RMT intensity (CSE100 and CSE120, respectively), in short-interval intracortical inhibition (SICI), and in intracortical facilitation (ICF). Each assessment comprised of ten trials for the pre, post1, and post2 measurement. Fig. 1 illustrates the sequence of assessments and tasks within each session. Here, we only report the results for the CSE measurements. Note that for the research question addressed here, the assessments of SICI and ICF were not analyzed.

**Figure 1:**
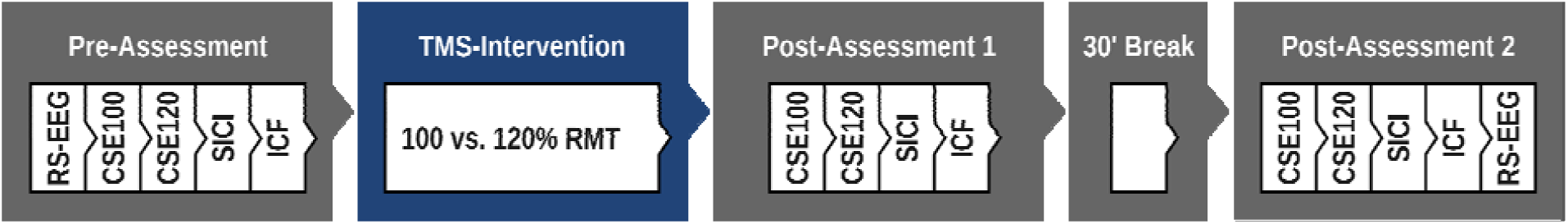
Sequence of tasks and assessments within each session. RS-EEG: resting state EEG; CSE100/120: corticospinal excitability, assessed at 100% or 120% of the resting-motor threshold (RMT); SICI: short-interval intracortical inhibition; ICF: intracortical facilitation.

### Electrophysiology

Electroencephalography (EEG) was recorded from 64 scalp channels using a BrainAmp amplifier (BrainProducts GmbH, Gilching, Germany) at a sampling rate of 1000 Hz and 16-bit resolution. We used a TMS-compatible cap that is compliant with the international 10-5 system (Easycap GmbH, Herrsching, Germany, online reference channel: FCz). Electrode impedances were kept below 5 kΩ. Surface electromyography (EMG) and electrocardiogram (ECG) were recorded from adhesive electrodes with Ag/AgCl contacts (Neuroline 720, Ambu A/S, Ballerup, Denmark). ECG was recorded in two-lead chest montage, EMG in bipolar belly-belly montage from the EDC, extensor carpi radialis (ECR), flexor carpi radialis (FCR), and flexor digitorum superficialis (FDS) muscles of both arms. Only data from the EDC, i.e., the target muscle for the experiment, are analyzed and reported here. In addition, we recorded breathing movements from a respiration belt (Vernier Software and Technology, Beaverton, USA; data not analyzed or reported here).

### TMS

For TMS, we used a triggerable MagPro X100 stimulator (MagVenture GmbH, Germany) with a figure-8 coil in combination with frameless stereotactic navigation (Localite GmbH, Germany).

### Hotspot search

The TMS hotspot was defined as the position of the TMS coil at which stimulation yielded the highest MEP amplitudes at the right EDC. To locate the hotspot, 40 bipolar TMS pulses at 45% of the maximum stimulator output (MSO) were applied at 40 distinct locations over the left hemisphere. The coil was held tangential to the head’s surface and was rotated 45° relative to the parasagittal plane. This is supposed to achieve optimal current flow and induce maximal MEP amplitudes [36]. Three additional pulses were applied at the three coil positions that had yielded the highest MEPs. The position with the highest mean amplitude was used as hotpsot for all subsequent TMS pulses.

### Single-pulse TM-intervention

TMS was triggered by a custom-built integrated measurement and stimulation device (IMSD; Neuroconn GmbH, Ilmenau, Germany). The IMSD provides a platform for categorizing brain states and controlling stimulation through real-time EEG analysis. During the experiment, the IMSD continuously measured EEG data from one bipolar channel (1cm anterior and posterior from C3) and extracted oscillatory phase at the target frequency across a 250 ms window by Discrete Fourier Transform (DFT) using the Goertzel algorithm [37]. The IMSD triggered TMS by a 5 V TTL pulse as soon as the EEG signal hit the predefined target phase at the target frequency. TMS pulses were applied at either 100% or 120% of the RMT, using biphasic pulses with a pulse width of 280 µs.

### Pre-/post TMS assessments of CSE

The TMS parameters for assessing pre-and post-intervention CSE were identical to those during the single-pulse TMS interventions. Ten single-pulses were applied at 100% or 120% of the RMT for each assessment. We quantified CSE within each assessment by calculating the median of the log-transformed MEPs of all 10 trials.

### Signal pre-processing

Signals were preprocessed and analyzed in Python, using the packages mne [38], mneiclabel [39,40], meegkit [41], pyprep [42], autoreject [43], antropy [44], and custom code.

### EEG data recorded during the TMS intervention

Raw EEG data were epoched from 3 s before to 3 after pulse onset. Then, the TMS pulse artifact was removed at each electrode using cubic interpolation of the raw EEG time-series in the time domain between-0.005 s and 0.015 s around the TMS-pulse [45].

Epochs were then cropped from-1.5 to 1.5 s around pulse onset. Next, we detected bad channels. To do so, we first concatenated the epoched data to a continuous dataset. Then, we calculated the standard deviation of the data of each channel from its spherical interpolation. Channels in which the actual signal deviated by more than 3 standard deviations from their interpolated signal for at least 40% of samples were marked as bad. We further identified bad channels using pyprep [42], which is the Python implementation of the PREP pipeline [46]. Channels were labeled as bad based on their signal-to-noise ratio, their overall amplitudes, the inter-channel correlations computed from 1s windows, and a random consensus approach using 1s data windows [43]. All bad channels were interpolated, and the EEG data were then re-referenced to the common average. Next, we applied notch-filters at 50 Hz (line noise) and all harmonic frequencies up to 250 Hz. We then performed robust polynomial detrending [47] with an order of 5 for each channel and each epoch using the package meegkit [41]. Finally, data were low-pass filtered at 40 Hz.

Eye-movement artifacts were removed using independent component analyses (ICA). First, we copied the original data and band-pass filtered the copied data between 1-30 Hz. Next, we automatically rejected bad data epochs using the package Autoreject with default parameters [43]. Then, an ICA was performed with correction for rank deficiency due to channel interpolation. Eye-movement related independent components were identified automatically using the MNE-ICALabel package [39,40] and were confirmed by visual inspection. These components were then rejected from the original data. In the cleaned EEG, bad epochs were identified and rejected using Autoreject (default settings), and the data were down-sampled to 500 Hz. Finally, data were surface Laplacian transformed.

### Resting-state EEG data

The raw data were notch filtered at 50 Hz and all harmonic frequencies up to 450 Hz and were then bandpass filtered between 0.5 and 40Hz. Then, the same pre-processing pipeline that had been used for the TMS-data were applied to the epoched resting-state EEG data (bad channel rejection, detrending, removal of eye-movement artifacts, rejection of bad epochs). Cleaned epochs were labelled with respect to the corresponding condition (eyes-open vs. eyes-closed). Finally, data were surface Laplacian transformed.

### Pre-computation of SVD entropy

SVD entropy was computed on the surface Laplacian transform of the EEG data. Entropy was calculated in the time domain separately for every electrode as suggested by Roberts et al. [26] and as implemented in the Python package antropy [44]. We used an embedding dimension of 3 samples and a delay of 1 sample. Entropy was normalized to one. We calculated entropy for a 1 s pre-pulse window ranging from-1.05 to-.05 s before pulse onset. For resting state EEG, entropy was extracted for every full 1 s epoch, and was then averaged across epochs within condition.

### Pre-processing of pre-pulse muscle tone

Raw EMG data from the target muscle were epoched from 3 s before to 3 s after pulse onset, and were then notch filtered at 50 Hz and all harmonic frequencies up to 200 Hz. Then, the time series for all trials were detrended with a 5th order polynomial using the robust detrending approach as implemented in meegkit [41].

To obtain an estimate of the pre-pulse muscle tone, data were cropped to the pre-pulse window (-1.05 to-0.05 s before pulse onset). Then, we calculated the log-transformed standard deviation of the time series separately for every epoch.

### Calculation of MEP amplitudes during the intervention

We determined MEP size as the peak-to-peak voltage fluctuation of the EMG signal over the EDC muscle in a time window of 15-60 ms after each TMS pulse. MEP sizes smaller than 50 µV were, by convention, considered subthreshold noise, and excluded from further analyses. Furthermore, we excluded all trials with an MEP size > 5000 µV, as well as all trials with technical flaws in the IMSD (e.g., no signal available for assessment of phase adherence). Overall, 19.5% of trials were excluded based on these criteria.

In some participants, more than 300 pulses had been applied during the intervention.

For consistency, only the first 300 pulses were kept for analysis.

### Calculation of CSE for the pre-and post-intervention assessments

Single-trial amplitudes of MEPs were calculated as described above. We computed the median MEP amplitude (after log-transforming) separately for each assessment (CSE100, CSE120), each time point (pre, post1, post2), and each session within each participant. Only data with at least 5 valid trials per measurement were included. For each session within each participant, we calculated the relative change in MEP amplitudes relative to the baseline (rel. change = (post-pre)/pre). A relative change of 0.10 thus reflects a 10% increase in MEP amplitude relative to the pre-intervention baseline.

### Data exclusion

For the analyses of the single-trial data obtained during the single-pulse TMS intervention, trials were rejected either if the EEG was highly artifact laden (see pre-processing pipeline), or if the log-transformed MEP amplitude exceeded more than 2 standard deviations from the mean within a specific session.

### Significance testing

Hypotheses were tested using linear mixed model analyses. Mixed models included random intercepts for participants and for sessions nested within participants. For all analyses, full models that included all relevant main effects and their interaction terms as fixed effects were fitted. All analyses were performed in R [48] but were called from within Python using the library pymer4 [49]. Linear mixed models were fitted using the R package lme4 [50]. T-values and corresponding p-values for all terms within a model were calculated the R-package lmertest [51]. Degrees of freedom for all models were estimated using the Satterthwaite method (default procedure in lmertest). For models that included categorical fixed effects containing more than two categories, type-III ANOVA summaries were obtained the R package lmertest (degrees of freedom estimation: Satterthwaite medhod).

If and when linear mixed models failed to converge, we dropped the random factor session (nested within participants). If mixed models still failed to converge, or mixed modeling was not advised (e.g., because the dependent variable was normalized within subject and session), linear models were fitted without random effects.

We ran separate models for each electrode and performed false-discovery rate (FDR) correction of p-values (Benjamini/Yekutieli method [52], FDR =.05) for multiple comparisons across electrodes (N=64) separately for each term in the model.

For all analyses, we normalized covariates to compute standardized β coefficients for each term in the fitted models. These coefficients represent the change in the dependent variable, measured in standard deviations, for each standard deviation change in the factor. As such, they provide a measure of effect size, allowing for direct comparison of the relative influence of different independent variables within the model.

## Results

### Pre-pulse entropy and TMS-induced MEPs

First, we investigated how pre-pulse entropy at each electrode accounts for the magnitude of TMS-induced MEPs at the single-trial level. To this end, we performed linear model analyses with normalized (within subject and session) log-transformed MEP amplitude as an outcome variable, and with fixed effects for stimulation intensity (effect coded: low-intensity =-1, high-intensity =+1), normalized pulse count, normalized pre-pulse muscle tone, normalized local pre-pulse entropy, and all interactions terms. No random effects were modeled because the dependent variable as well as all independent variables (pulse count, muscle tone, and entropy) were normalized within participant and session. We ran separate models for each electrode and performed false-discovery rate correction of p-values [52] for multiple comparisons across electrodes (N=64) separately for each term in the model.

### Factors influencing MEP amplitude

Pulse count and pre-pulse muscle tone were included as independent variables in the models because both factors significantly accounted for the variability in MEP amplitudes (see Supplementary Analysis 1). Specifically, MEP amplitudes were higher when pulses were applied during states of elevated muscle tone. This is in line with previous reports suggesting that muscle activation significantly accounts for variability in MEP amplitudes [53]. MEP amplitudes further generally increased with the number of pulses applied. By controlling for these factors, we ensured that any observed effects of entropy were not due to other factors.

High pre-pulse entropy over right fronto-central electrodes was associated with larger MEP amplitudes after controlling for all other effects (Fig. 2A). An overview of all effects at every electrode is provided in Fig. S1. High entropy over frontal electrodes as well as over left central electrodes, i.e. the site of stimulation, further amplified the effect of pre-pulse muscle tone on MEP amplitudes. This was suggested by the significant interaction (Fig. 2A) between entropy and muscle tone (MEP∼E:M). Importantly, a significant four-way interaction between intensity, entropy, pulse count, and muscle tone (MEP∼I:E:P:M) suggested that the influence of entropy was not uniform across stimulation intensity.

**Figure 2:**
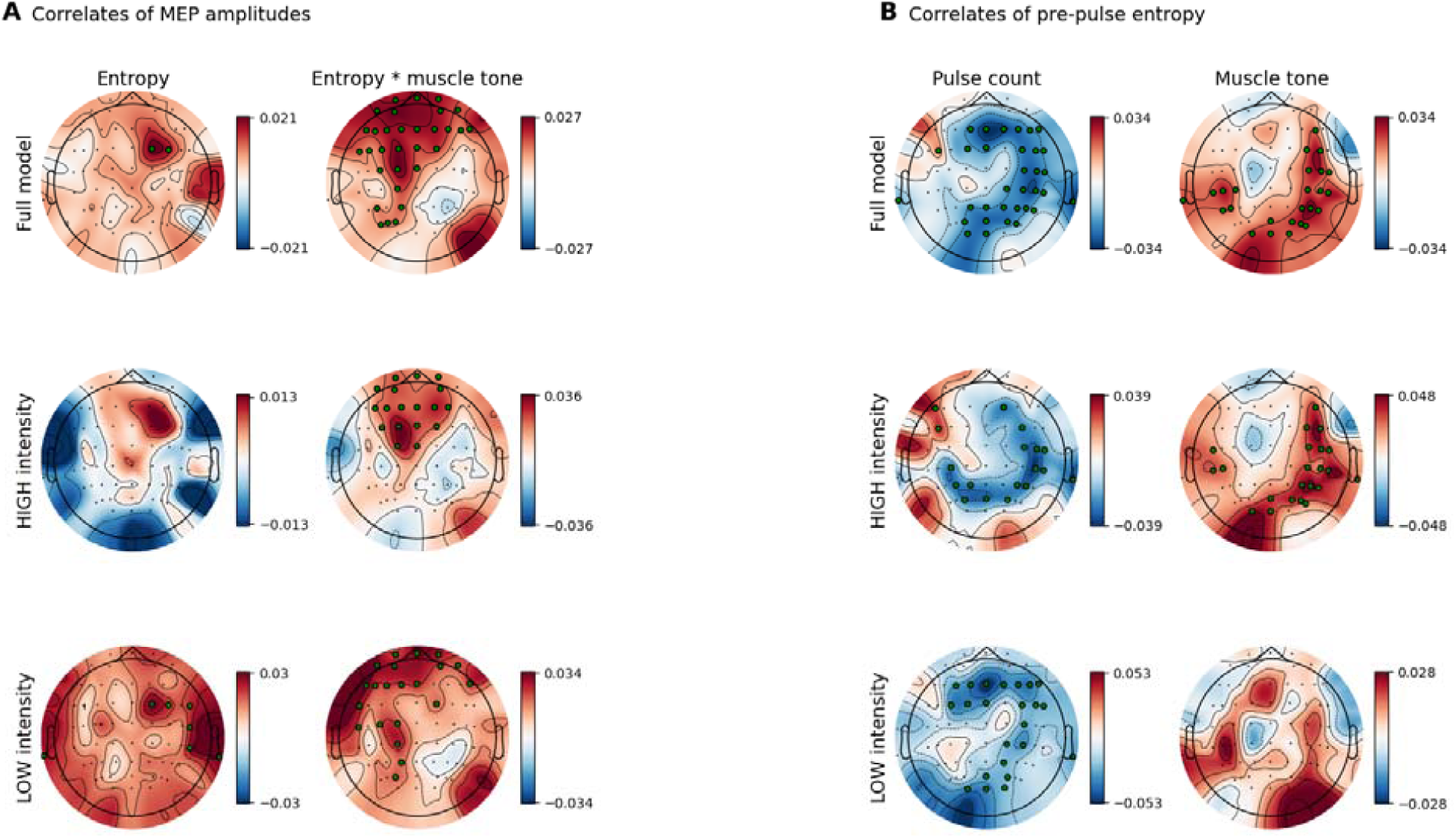
**Correlates of MEP amplitudes (A) and pre-pulse entropy (B) throughout the intervention**. A) pre-pulse entropy (E) as well as the interaction term between E and pre-pulse muscle tone (E*M) were significantly associated with MEP amplitudes in the full model that included both the high and low intensity intervention. When tested separately for both interventions, E*M remained significant in both models, but E alone was associated with MEP amplitudes only at low stimulation intensity. B) pre-pulse entropy (E) decreased on most electrodes as function of pulse counts (P). This was observed in the full model that included both interventions (high & low), as well as in separate models for both interventions. Elevated pre-pulse muscle tone was further associated with heightened entropy over right-hemispheric electrodes, but only for the high intensity intervention; topoplots provide the standardized β coefficients for a given term from the statistical models for each electrode. Significant electrodes (FDR corrected for multiple comparisons) are highlighted.

### Effect of stimulation intensity on entropy-MEP relationship

To elucidate the differential effects of pre-pulse entropy on MEP amplitudes at distinct stimulation intensities, we performed separate analyses for the high-vs. low-intensity stimulation interventions. In both interventions, the interaction between high frontal as well as left central entropy and muscle tone remained significant (see Fig. 2A). The interaction suggests that states of elevated excitability in the motor cortex boosted the impact of TMS pulses on MEP amplitudes if the corresponding muscle was tone was elevated. The interaction between entropy and muscle tone appeared to be less prominent (fewer significant electrodes) at high stimulation intensity. Hence, at high stimulation intensity, the current state of cortical excitability had a smaller impact on the magnitude of MEP amplitudes due to the intensity of the stimulation pulse.

At low stimulation intensity, elevated right frontal entropy alone was associated with higher MEP amplitudes (see Fig. 2A). This effect was also present at high intensity, but was weaker (low: β = 0.030 vs. high: β = 0.013) and did not reach significance. This indicates that at high stimulation intensity, the current state of cortical excitability had a smaller impact on the magnitude of MEP amplitudes.

### Changes in entropy during and after the intervention

We further investigated how stimulation altered entropy during the intervention and the entropy from before to after the interventions. Pre-pulse entropy significantly decreased throughout the stimulation on most electrodes, except those located over the left motor cortex (Fig. 2B). This was suggested by linear model analyses with normalized (within subject and session) entropy as an outcome variable, and with fixed effects for stimulation intensity (effect coded: low-intensity =-1, high-intensity =+1), normalized pulse count, and normalized pre-pulse muscle tone. The analysis further revealed that entropy was significantly elevated when muscle tone was high (Fig. S2). These effects were highly similar across stimulation intensities. However, the association between muscle tone and entropy did not reach significance in the low intensity stimulation condition.

Entropy as measured during resting did not significantly change from before to after the interventions. This was suggested by linear mixed models in which we analyzed the impact condition (eyes open vs. eyes closed; effect coded), time (pre vs. post intervention; effect coded) and the type of intervention (high vs. low intensity; effect coded) on entropy (normalized across all participant and session) at each electrode (random intercepts for participants and sessions nested within participants). Only the factor of eyes open vs. eyes closed reached significance (Fig. S5). In fact, entropy was significantly higher during eyes open as compared with eyes closed at most electrodes (Fig. S5). This difference was most prominent at occipital and parietal electrodes (β = 0.517).

### Entropy as a marker of increased excitability

Although the interventions did, on average, not lead to a change in resting-state entropy from before to after the interventions, participants who showed a larger increase in entropy close to the stimulation site also displayed larger relative changes in corticospinal activity. Specifically, an increase in resting state entropy close to the stimulation site from before to after the high-intensity intervention measured during both, eyes open and eyes closed conditions was associated with a larger relative change in corticospinal activity measured at 100% intensity (CSE100) during the post1 and the post2 assessments (Fig. 3; Fig. S4; β > 0.47). This was observed in linear mixed models where the relative change for each assessment (CSE100, CSE120; normalized across participants and session) was used to identify associations with the change in entropy (post minus pre; normalized across participants and sessions). Models were run separately for each assessment, each time point (post1 and post2), each intervention (high vs. low intensity), and each resting state condition (eyes open vs. eyes closed). See Fig. S4 for an overview of all results. Hence, participants who showed a larger relative increase in corticospinal excitability (measured at 100% intensity) induced by the high-intensity intervention also showed a higher increase in entropy. In sum, this suggests that entropy is a marker of increased excitability.

**Figure 3:**
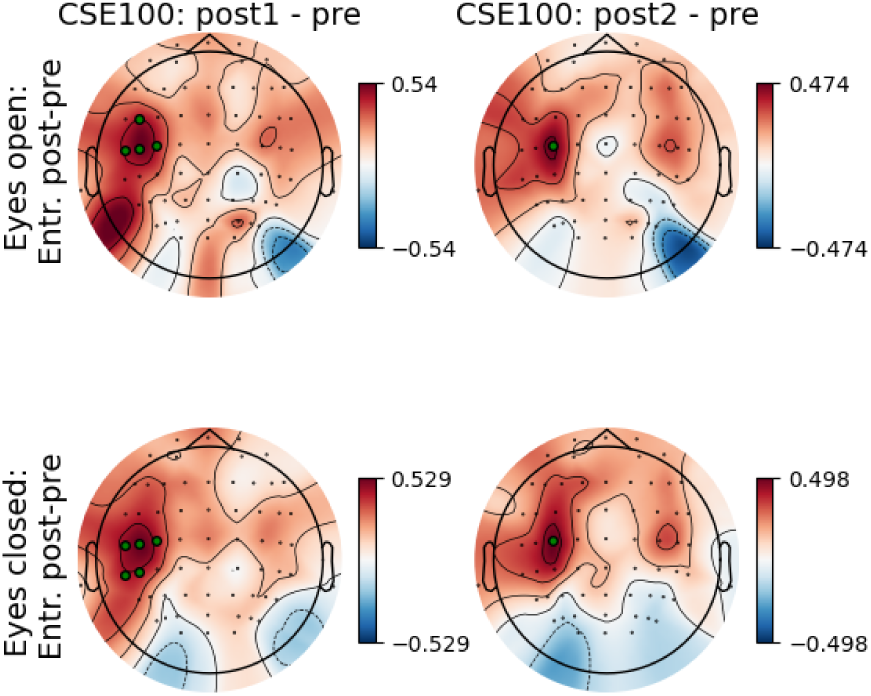
Associations between pre-post changes in MEP amplitude and entropy. A larger increase in resting state entropy from pre-to post-intervention was associated with a larger increase in corticospinal excitability measured at 100% intensity (CSE100). This association was present only following the high-intensity intervention (plotted here) and was observed close to the stimulation site both in the eyes open and eyes closed resting condition and for the post1 and post2 assessment; topoplots provide the standardized β coefficients for a given term from the statistical models for each electrode. Significant electrodes (FDR-corrected) are highlighted (green dots).

Importantly, resting state entropy and corticospinal excitability (CSE100) were not measured simultaneously, but in sequence. The observed association between entropy and excitability thus reflect lasting excitability changes induced by the intervention at high stimulation intensity.

## Discussion

Entropy – often conceptualized, informally, as “complexity” [54], the unpredictability of a signal, or the number of ways in which a signal can be arranged – has found a number of applications in recent years as a marker of sleep state [55–57], conscious state [54], and even diagnosis or prognosis in certain brain disorders [21,58–60]. Here, instead of relating entropy to relatively stable brain states which change over the course of hours or even years, we related entropy to rapid fluctuations in CSE, building on findings from a recent EEG study on the association between pre-pulse entropy and MEPs following randomly applied single TMS pulses [25] and focusing in the present study on the evolution of entropy and CSE during and after an intervention of 300 single TMS pulses.

Given that low neural entropy recorded from cortical activity is generally associated with low or no responsiveness to afferent signals (e.g., during deep sleep or general anesthesia), we expected that states of higher cortical entropy would be more receptive to electromagnetic stimuli delivered via TMS. We indeed found evidence to support this hypothesis. Notably, while a number of studies [61–64] have looked at the entropy of cortical activity after a TMS pulse (i.e., perturbational complexity [61]) as a readout of conscious state, our study continues a novel approach of measuring neural entropy before a TMS pulse, specifically, as a correlate of CSE [25].

Specifically, we found that right frontal pre-pulse entropy related positively to CSE, as measured by the amplitude of the resulting MEP. The right lateralization of electrodes showing a significant effect can be seen in Fig. 2A both for the “full” and “low intensity” entropy models of MEP amplitudes. A qualitative similar, albeit weaker and non-significant pattern was observed in the “high intensity” model. The right lateralization of this effect is somewhat surprising, given that we stimulated left motor cortex. This could be explained by interhemispheric inhibition [65], in which the right frontal cortex regulates the excitability of the left motor cortex.

Interestingly, this location where entropy was associated with MEP amplitudes overlaps with the topography of the late component of TMS-evoked EEG potentials (TEPs). Single-pulse TMS over the left motor cortex induces TEPs characterized by a late frontal and contralateral positivity (e.g. [66,67]). This late component is thought to serve as a global marker of cortical excitability [68]. Indeed, pharmacological agents that reduce neural excitability have been shown to attenuate the amplitude of this TEP component [67,69]. The topographic similarity between the observed pre-pulse entropy distribution and TEP maps associated with cortical excitability further suggests that elevated entropy over right frontal electrodes reflects increased cortical excitability in the neural network facilitating corticospinal propagation of TMS-induced activity.

Entropy was most consistently associated with MEPs in interaction with muscle tone across electrodes. This interaction was observed over frontal and central electrodes, with a tendency for a left lateralization (Fig. 2A). This suggests that high fronto-central excitability as indicated by elevated entropy mediated the cortical and corticospinal propagation of TMS-induced activity, but that the impact on MEP amplitudes was contingent on an elevated tone of the target muscle. The fact that entropy was not directly associated with pre-pulse muscle tone at the relevant electrodes (Fig. 2B) suggests that entropy did not merely mirror elevated muscle tone or cortical processes that moderate muscle tone. Interestingly, the interaction between entropy and muscle tone showed similar magnitudes and similar topographic distributions at both stimulation intensities supporting the robustness of the finding. However, entropy overall appeared to be associated with MEP amplitudes more strongly at low intensity stimulation. This is in line with previous work suggesting higher brain state dependence of TMS-induced responses at low stimulation intensities [70,71].

Interestingly, entropy not only predicted the impact of TMS pulses, but was itself modulated by the TMS interventions. This was suggested by the finding that the global decrease in entropy throughout the interventions was absent at the site of stimulation. In the high intensity intervention, entropy even increased in left frontal electrodes (Fig. S2 B). The global decrease of entropy might have occurred independently of stimulation and could be a mere consequence of increasing fatigue. Previous studies suggested that fatigue [72–74] or relaxation [75] lower EEG entropy, possibly reflecting reduced vigilance during prolonged TMS interventions [56]. Importantly, TMS appeared to counteract, disrupt, or prevent this decrease at the site of stimulation, especially during high-intensity stimulation. This suggests that the changes induced by TMS pulses lead to a local increase in neural excitability and hence entropy that counteracted the overall decrease in entropy. Additionally, a local increase in entropy from the pre-to the post-intervention resting-state EEG was associated with an increased corticospinal excitability from the pre-to the post intervention assessments. During the pre-and post-intervention resting-state EEG recordings, no TMS coil was used, confirming that the observed entropy changes were not artifacts caused by coil contact with the electrode. Therefore, entropy mirrored corticospinal excitability both during and after the intervention.

The variability in motor excitability that was accounted for by entropy was relatively low (standardized β values <.040) when compared to other variables such as the number of pulses applied (β > 0.047) and particularly pre-pulse muscle tone (β > 0.57). Despite participants being instructed to keep their hand fully relaxed, subtle involuntary muscle activity may have persisted, contributing to elevated baseline muscle tone. Such low-level tonic EMG activity, often undetectable without continuous monitoring, apparently influenced corticospinal excitability. Furthermore, the use of the psychomotor vigilance task during the intervention, interspersed pseudo-randomly between TMS pulses to maintain alertness, may have further influenced motor system excitability. The requirement to fixate on a grid of red LEDs and respond rapidly with the left index finger likely increased general arousal and attentional engagement, potentially reducing intracortical inhibition and enhancing baseline excitability, while also elevating muscle tone in the targeted right finger extensor muscles. This highlights the difficulty of finding specific cortical markers of elevated corticospinal excitability that are independent of these confounding factors, as is also the case with EEG power [76] or phase [77], or their interactions [78]. Moreover, this emphasizes the importance of taking into account extracortical covariates such as muscle tone for modeling and describing fluctuations in corticospinal excitability.

While our findings suggest that entropy measures could serve as novel markers for optimizing brain state-dependent TMS applications, our study does not address whether selectively targeting high-entropy states leads to greater excitability changes.

Interestingly, the association between TMS-induced changes in entropy and induced changes in CSE (pre-post measurements) was large (β > 0.47), unlike the association between pre-pulse entropy and CSE during the intervention. This finding can be interpreted in several, not mutually exclusive, ways. One possibility is that the association developed over time through mechanisms such as LTP-like effects, changes in connectivity, or shifts in the excitation/inhibition balance. This would suggest that entropy reflects a brain state conducive to plasticity or network reconfiguration – processes that emerge over time rather than immediately. Alternatively, task-related increases in muscle tone during the intervention may have confounded the pure effect of entropy, which became more apparent post-intervention when the task was no longer being performed. In any case, these findings suggest that entropy can be used as marker of the TMS-induced changes in cortical excitability. This is supported by a recent study by Liu et al. [79] in which the authors observed an increase in fMRI-derived entropy following a high-(120% RMT) but not a low-intensity (90% RMT) intermittent theta burst TMS intervention.

### Could entropy serve as general measure of cortical activity that extends to other tasks?

Similar to the variability of TMS-probed cortical excitability at multiple timescales is the diurnal variation in alertness, attention, and performance in cognitive and sensorimotor tasks [6,80]. For example, the ability to maintain attention is impaired by sleep deprivation, displays systematic circadian variation [81], and is affected by the presence of spontaneous slow oscillations in the electroencephalogram (EEG) that reflect brief intrusions of sleep-like activity during wakefulness [82]. Hence, both the brain’s susceptibility to magnetic stimulation and cognitive performance depend on the current state of cortical excitability [83,84]. Not only global brain-wide states [82], but also states within specific task-related neuronal networks [82,85] may shape cognitive performance and the brain’s response to stimulation [7,81,86]. Entropy as a marker of cortical excitability might thus also be useful for mapping fluctuations in cognitive performance. Such a marker might further inspire new applications for improving our day-to-day life. For example, it might allow for measuring a driver’s alertness during a long drive, or for assessing students’ vigilance in the classroom.

### Limitations, future directions, and conclusions

Despite its promising findings, this study has several limitations. First, the study focused on a single-pulse TMS intervention, limiting generalizability to repetitive or patterned stimulation paradigms [79]. Second, while entropy provides valuable insights, its relationship with other EEG-derived measures of excitability, such as power spectral density or connectivity metrics, remains unclear. Third, the study design did not directly assess causal mechanisms linking entropy to corticospinal excitability, leaving open questions about underlying neurophysiological processes. Finally, the complex interactions between stimulation intensity, muscle tone, pulse count and entropy necessitate further investigation and larger sample sizes to improve reproducibility and to disentangle online and offline effects – both in the presence and absence of concurrent tasks that affect alertness, attention, cognition, and muscle tone.

Future research should extend the present findings by investigating whether entropy is associated with responses to different TMS paradigms, e.g., including repetitive or theta burst stimulation [79]. Additionally, studies should explore the neurobiological underpinnings of entropy’s role in excitability, potentially using multimodal approaches. Examining how entropy-driven brain state-dependent TMS influences excitability is the natural next step and could advance personalized neuromodulation interventions. Lastly, refining entropy-based biomarkers across different populations and experimental conditions will be critical for establishing their translational relevance in both research and clinical applications.

In conclusion, this study provides evidence that entropy, derived from singular value decomposition of 1s segments of electroencephalography measurements, serves as a marker of cortical excitability and its fluctuations. Higher pre-pulse entropy, particularly over frontal electrodes, is associated with increased motor-evoked potentials, indicating a link between ongoing neural complexity and corticospinal excitability. Furthermore, TMS appears to locally enhance entropy at the stimulation site, while entropy is decreasing elsewhere, suggesting a regionally specific interaction between brain state and stimulation effects. Importantly, participants who exhibited greater post-intervention increases in entropy near the stimulation site also showed larger corticospinal excitability changes, reinforcing the promise of entropy as a biomarker for modulated excitability. These findings highlight the potential of entropy for optimizing intervention protocols by informing brain state-dependent stimulation strategies and evaluating intervention effects.

## CRediT authorship contribution statement

Simon Ruch: Conceptualization, Methodology, Formal analysis, Visualization, Writing - Original draft.

Joel Frohlich: Conceptualization, Methodology, Formal analysis, Writing - Review & Editing.

Marius Keute: Data curation, Methodology, Validation, Writing - Review & Editing. Ge Tang: Investigation, Data curation, Writing - Review & Editing.

Nelli Keksel: Investigation, Data curation, Writing - Review & Editing.

Alireza Gharabaghi: Conceptualization, Resources, Supervision, Project administration, Funding acquisition, Writing - Review & Editing.

The authors have read and approved the manuscript, which has undergone thorough revisions. Additionally, the manuscript was refined for grammatical accuracy, sentence structure and wording using a convolutional neural network-based tool (DeepL) and an advanced language model (ChatGPT).

## Data and code availability

The preprocessed data (precomputed entropy values, precomputed muscle tone, amplitudes of motor evoked potentials) and the code and output supporting the key findings of this study are available on OSF: https://doi.org/10.17605/OSF.IO/GAMVR. Additional data will be made available for researchers upon request by the first author after journal publication, in accordance with ethical and legal regulations.

## Acknowledgments

We warmly thank all participants who volunteered for our study. We also gratefully acknowledge support from the Open Access Publishing Fund of the University of Tübingen.

Simon Ruch is currently affiliated with UniDistance Suisse, Faculty of Psychology, Brig, Switzerland. Joel Frohlich is currently affiliated with fMEG Center, University of Tübingen, Tübingen, Germany.

## Supplementary material

### Supplementary analysis 1: The impact of pre-pulse muscle tone and number of applied pulses on single-pulse MEPs during the intervention

To assess the main determinants of the temporal variability of MEP amplitudes during the interventions, we performed a linear mixed model analysis with log-transformed MEP-amplitudes as dependent variable and fixed effects for stimulation intensity (effect coded: low-intensity =-1, high-intensity =+1), normalized pulse count, normalized pre-pulse muscle tone, and all interactions terms. We modeled random intercepts for participants and sessions nested within participants.

The analysis confirmed that amplitudes of the motor evoked potentials were significantly larger in the high-intensity intervention (main effect of intervention: β = 0.727, t(118.007) = 25.649, p <.001). Importantly, amplitudes were significantly larger when pulses were applied during periods of elevated muscle tone (main effect of muscle tone: β = 0.273, t(33678.034) = 106.641, p <.001). The MEP amplitudes further increased throughout the intervention (main effect of pulse count: β = 0.029, t(33678.035) = 11.491, p <.001). Significant interactions between intensity and muscle tone (β =-0.019, t(33678.035),-7.319, p <.001) as well as between intensity and pulse count (β =-0.025, t(33678.034) =-9.776, p <.001) suggested that both, the effect of muscle tone and of the number of pulses applied were less prominent in the high-intensity intervention. A significant negative interaction term between pulse count and muscle tone (β =-0.009, t(33706.898) =-3.383, p =.001) further indicated that the impact of muscle tone on MEP amplitudes dissipated over time. This tendency was similar across interventions (non-significant three-way interaction between stimulation intensity, muscle tone, and pulse count: p =.913).

### Supplementary analysis 2: The impact of stimulation intensity and pulse count on pre-pulse muscle tone during the interventions

Pre-pulse muscle tone was overall higher during the high-intensity intervention (main effect of intervention: β = 0.325, t(117.957) = 7.677, p <.001), and increased significantly throughout the intervention (main effect of pulse count: β = 0.030, t(33681.937) = 7.257, p < 0.001). This was suggested by a linear mixed model analysis with normalized muscle tone as dependent variable, intensity, normalized pulse count, and their interaction term as fixed effects, and random intercepts for participants and sessions nested within participants. Importantly, a significant interaction between pulse count and stimulation intensity (β = 0.041, t(33681.937) = 9.869, p <.001) suggested that the effect of stimulation was not uniform across interventions. Post-hoc tests revealed that muscle tone increased with pulse counts at high intensity stimulation (120%: β = 0.069, t(16674.003) = 11.763, p(uncorrected) <.001), but rather tended to decrease at low-intensity stimulation (100%: β =-0.012, t(17007.849) =-1.903, p(uncorrected) =.057).

### Supplementary analysis 3: Long-term impact of single-pulse interventions on corticospinal excitability

Prior to each intervention (pre), immediately thereafter (post1), and after a delay of 30 minutes (post2), we administered standardized protocols to assess corticospinal excitability (probed at 100% and 120% intensity, CSE100 and CSE120, respectively).

To assess the long-term impact of the TMS interventions on corticospinal excitability, we performed linear mixed model analyses with the relative change in CSE as outcome variable (normalized across participants), and with time (ordered factor with the constant pre-value = 1 and the relative change for post1 and post2) and intervention (high vs. low intensity) as independent variables. Separate analyses were performed for the assessment of CSE100 and CSE120.

Corticospinal excitability measured at 100% RMT intensity (CSE100) increased significantly from before to after the interventions (main effect of time: F(2, 385.078) = 9.468, p <.001). Descriptively, the relative change in corticospinal excitability was largest (>10% increase) in the post1 measurement following the high-intensity intervention (see Fig. S3, left panel). However, neither the effect of intensity (F(1, 385.078) = 1.470, p =.226) nor the interaction between intensity and time (F(2, 385.078) = 0.779, p =.460) reached significance.

On average, corticospinal excitability did not change if measured at 120% intensity (CSE120; no main effect of time: F(2, 384.010) = 0.548, p =.579). However, a significant main effect of intensity (F(1, 384.017) = 17.122, p <.001) and the significant interaction between time and intensity (F(2, 384.017) = 4.319, p =.014) suggested that high-vs. low-intensity interventions affected corticospinal excitability differently when probed at 120% intensity. Post-hoc tests revealed that excitability increased only following the high-(significant main effect of time: F(2, 184.106) = 4.124, p(uncorrected) =.018) but not the low-intensity intervention (no main effect of time: F(2, 184.004) = 0.919, p(uncorrected) =.401).

## Supplementary figures

**Figure S1:**
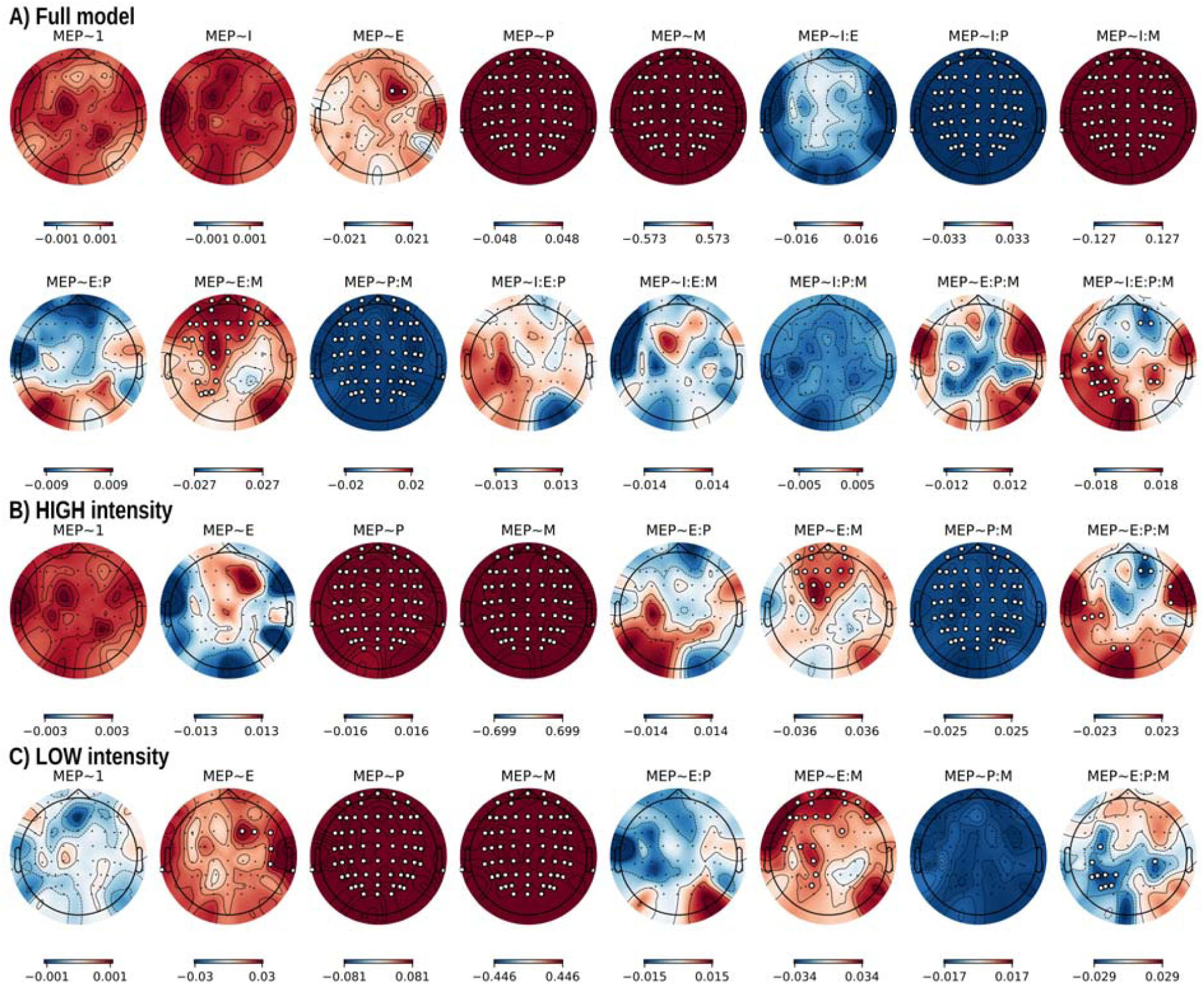
overview of all analyzed predictors of MEP amplitudes. Overview of the predictive value of al included predictors on motor-evoked potentials (MEP). A) Analysis of the full model combining both interventions (HIGH & LOW intensity). B) and C) separate models for the HIGH and LOW intensity interventions, respectively. Predictors are indicated in the title of each topographic map (MEP∼predictor): intercept (1), intensity (I) of the intervention (omitted in panel B and C), entropy (E), pulse count (P), muscle tone (M), and all possible interaction terms (:). Topographic maps display the standardized β coefficients obtained from each model calculated for each channel. Channels at which a predictor reached significance (after false-discovery rate correction for N=64 channels) are highlighted.

**Figure S2:**
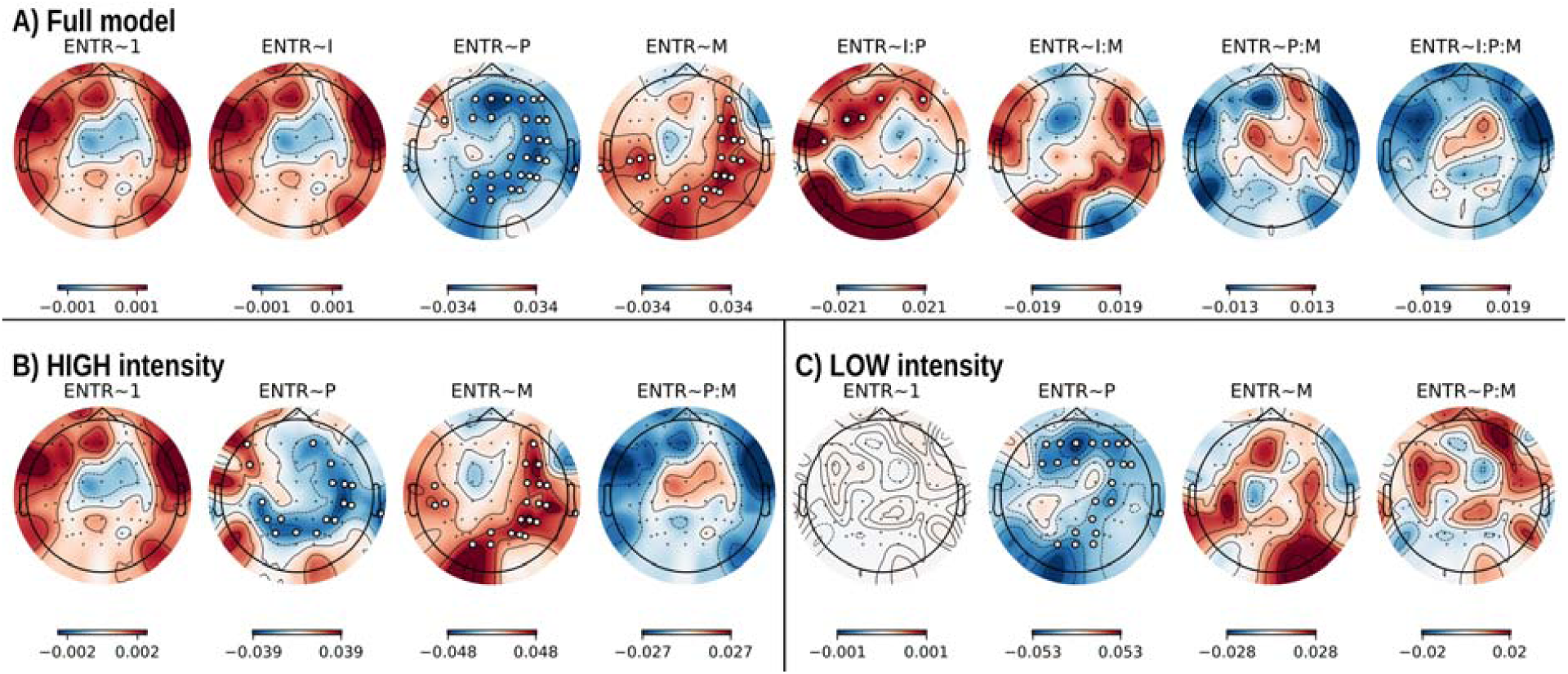
overview of all analyzed predictors of pre-pulse entropy. Overview of the predictive value of all included predictors on pre-pulse entropy (ENTR). A) Analysis of the full model combining both interventions (HIGH & LOW intensity). B) and C) separate models for the HIGH and LOW intensity interventions, respectively. Predictors are indicated in the title of each topographic map (ENTR∼predictor): intercept (1), intensity (I) of the intervention (omitted in panel B and C), pulse count (P), muscle tone (M), and all possible interaction terms (:). Topographic maps display the standardized β coefficients obtained from each model calculated for each channel. Channels at which a predictor reached significance (after false-discovery rate correction for N=64 channels) are highlighted.

**Figure S3:**
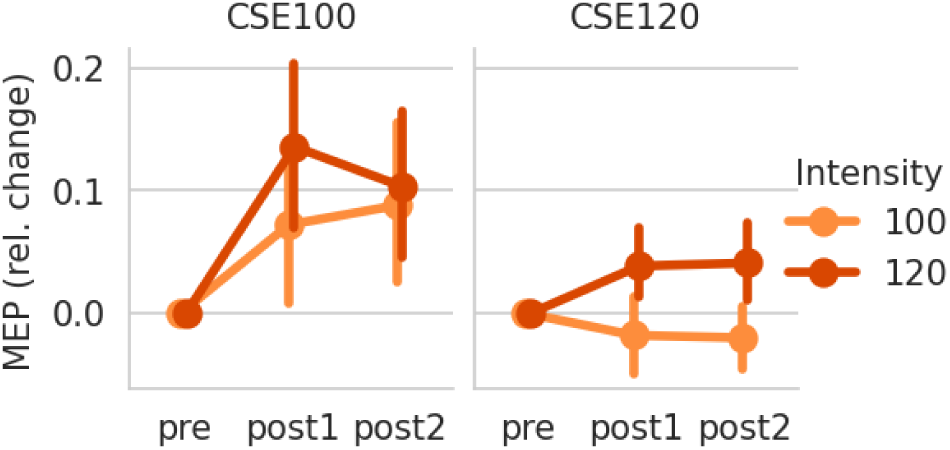
Pre to post changes in MEP amplitude. Relative change in corticospinal excitability (CSE100/120 measured at 100% and 120% RMT intensity) from pre, to post1, and post 2 measurement. Mean of relative change (post minus pre, divided by pre) with 95% confidence intervals is plotted. Relative change is plotted separately for the LOW (100%, in orange) and HIGH (120%, in red) intensity intervention.

**Figure S4:**
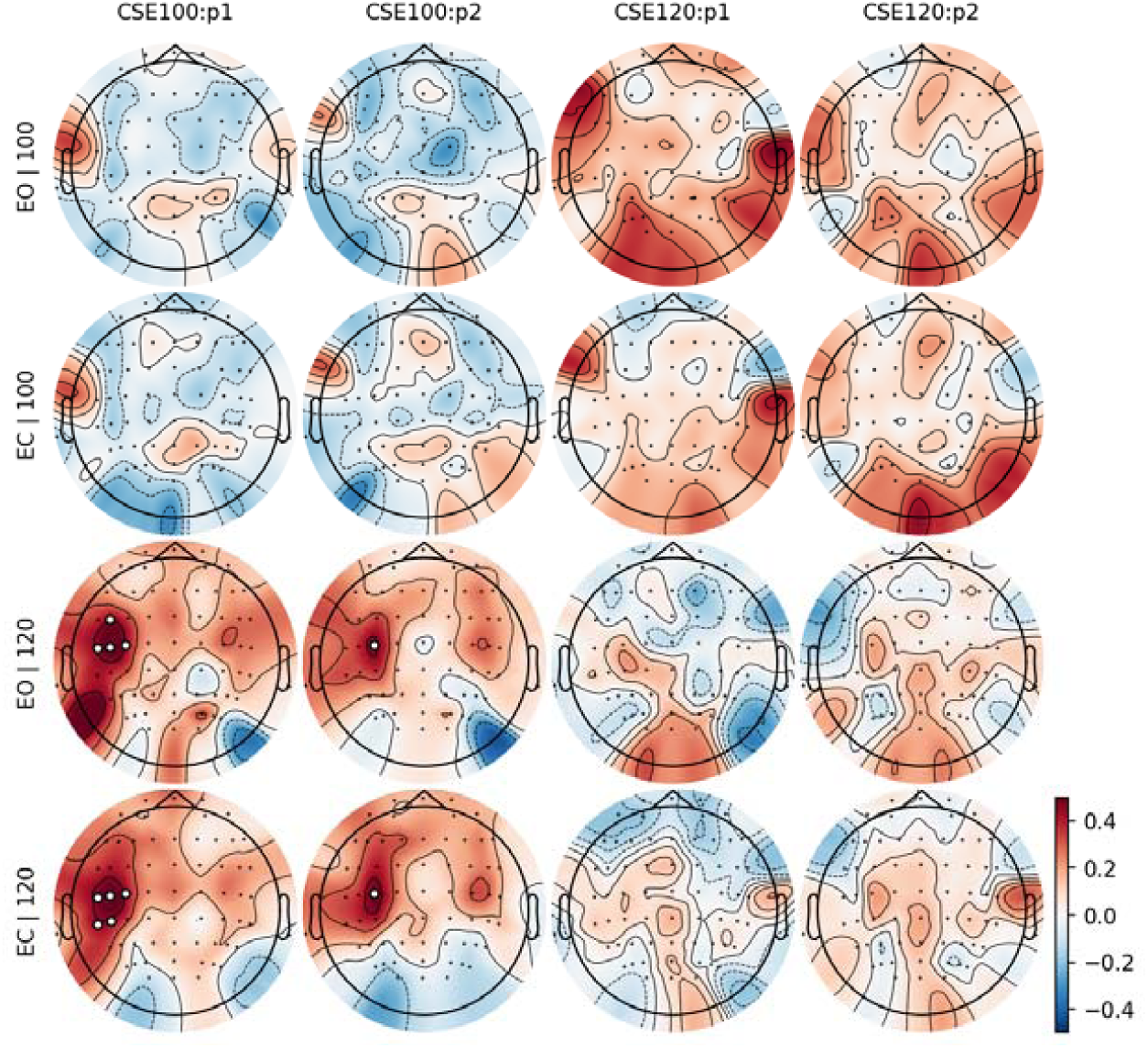
Associations between pre-post changes in MEP amplitude and entropy. Association between change in resting-state entropy and change in each of the TMS assessments (CSE100, CSE120) from before to after the intervention. All tests were performed separately for the different types of interventions (100% vs. 120% stimulation intensity), the different resting state conditions (EO: eyes-open, EC: eyes-closed), and for all assessments at the post1 and post2 time-point. In the 120% intensity intervention (third and fourth row), a significant association between the increase in excitability as measured with the CSE100 protocol (first and second column), and an increase in entropy was observed close to the stimulation site. Topographic maps display the standardized β coefficients obtained from each model calculated for each channel. Channels at which a predictor reached significance (after false-discovery rate correction for N=64 channels) are highlighted.

**Figure S5:**
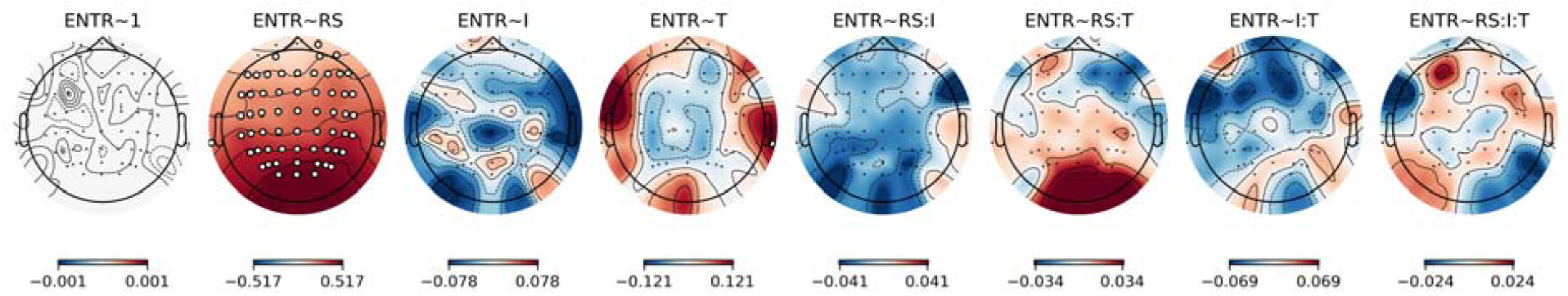
Pre-post changes in resting state entropy. Impact of resting state condition (RS: eyes open vs. eyes closed), high vs. low intensity intervention (I), time of testing (T: pre vs. post intervention) and all their interactions on entropy (ENTR) measured at rest. Entropy was significantly elevated in most electrodes in the eyes open compared to the eyes closed condition. This was independent of stimulation intensity (I) or time of testing (T; no other main effects or interaction terms reached significance). Topographic maps display the standardized β coefficients obtained from each model calculated for each channel. Channels at which an independent variable was significantly associated with entropy (after false-discovery rate correction for N=64 channels) are highlighted.

## References

[1] Bergmann TO, Mölle M, Schmidt MA, Lindner C, Marshall L, Born J, et al. EEG-guided transcranial magnetic stimulation reveals rapid shifts in motor cortical excitability during the human sleep slow oscillation. J Neurosci 2012;32:243–53. 10.1523/JNEUROSCI.4792-11.2012.

[2] Janssens SEW, Sack AT. Spontaneous fluctuations in oscillatory brain state cause differences in transcranial magnetic stimulation effects within and between individuals. Front Hum Neurosci 2021;15.

[3] Keute M, Gebühr J-S, Guggenberger R, Trunk BH, Gharabaghi A. Phase-specific stimulation reveals consistent sinusoidal modulation of human corticospinal excitability along the oscillatory beta cycle 2023:2023.04.25.538229. 10.1101/2023.04.25.538229.

[4] Schilberg L, Oever ST, Schuhmann T, Sack AT. Phase and power modulations on the amplitude of TMS-induced motor evoked potentials. PLOS ONE 2021;16:e0255815. 10.1371/journal.pone.0255815.

[5] Wischnewski M, Haigh ZJ, Shirinpour S, Alekseichuk I, Opitz A. The phase of sensorimotor mu and beta oscillations has the opposite effect on corticospinal excitability. Brain Stimulat 2022;15:1093–100. 10.1016/j.brs.2022.08.005.

[6] Ly JQM, Gaggioni G, Chellappa SL, Papachilleos S, Brzozowski A, Borsu C, et al. Circadian regulation of human cortical excitability. Nat Commun 2016;7:11828. 10.1038/ncomms11828.

[7] Salehinejad MA, Wischnewski M, Ghanavati E, Mosayebi-Samani M, Kuo M-F, Nitsche MA. Cognitive functions and underlying parameters of human brain physiology are associated with chronotype. Nat Commun 2021;12:4672. 10.1038/s41467-021-24885-0.

[8] Badawy RAB, Vogrin SJ, Lai A, Cook MJ. Cortical excitability changes correlate with fluctuations in glucose levels in patients with epilepsy. Epilepsy Behav 2013;27:455–60. 10.1016/j.yebeh.2013.03.015.

[9] Walker CP, McFerren AL, Rammani DR, Buse JB, Frohlich F. Metabolic state and gustatory perception shapes dynamic interplay between cortical excitability and motor response. Brain Stimul Basic Transl Clin Res Neuromodulation 2021;14:202–5. 10.1016/j.brs.2020.12.002.

[10] Civardi C, Boccagni C, Vicentini R, Bolamperti L, Tarletti R, Varrasi C, et al. Cortical excitability and sleep deprivation: a transcranial magnetic stimulation study. J Neurol Neurosurg Psychiatry 2001;71:809–12. 10.1136/jnnp.71.6.809.

[11] Manganotti P, Palermo A, Patuzzo S, Zanette G, Fiaschi A. Decrease in motor cortical excitability in human subjects after sleep deprivation. Neurosci Lett 2001;304:153–6. 10.1016/S0304-3940(01)01783-9.

[12] Haxel L, Belardinelli P, Ermolova M, Humaidan D, Macke JH, Ziemann U. Decoding motor excitability in tms using eeg-features: an exploratory machine learning approach 2024:2024.02.27.582361. 10.1101/2024.02.27.582361.

[13] Gharabaghi A, Kraus D, Leao MT, Spüler M, Walter A, Bogdan M, et al. Coupling brain-machine interfaces with cortical stimulation for brain-state dependent stimulation: enhancing motor cortex excitability for neurorehabilitation. Front Hum Neurosci 2014;8. 10.3389/fnhum.2014.00122.

[14] Lieb A, Zrenner B, Zrenner C, Kozák G, Martus P, Grefkes C, et al. Brain□oscillation-synchronized stimulation to enhance motor recovery in early subacute stroke: a randomized controlled double□blind three□ arm parallel□group exploratory trial comparing personalized, non□ personalized and sham repetitive transcranial magnetic stimulation (Acronym: BOSS-STROKE). BMC Neurol 2023;23:204. 10.1186/s12883-023-03235-1.

[15] Nagy B, Protzner AB, van der Wijk G, Wang H, Cortese F, Czigler I, et al. The modulatory effect of adaptive task-switching training on resting-state neural network dynamics in younger and older adults. Sci Rep 2022;12:9541. 10.1038/s41598-022-13708-x.

[16] Ahmad J, Ellis C, Leech R, Voytek B, Garces P, Jones E, et al. From mechanisms to markers: novel noninvasive EEG proxy markers of the neural excitation and inhibition system in humans. Transl Psychiatry 2022;12:1–12. 10.1038/s41398-022-02218-z.

[17] Flood MW, Grimm B. EntropyHub: An open-source toolkit for entropic time series analysis. PLOS ONE 2021;16:e0259448. 10.1371/journal.pone.0259448.

[18] Kostic MM. The elusive nature of entropy and its physical meaning. Entropy 2014;16:953–67. 10.3390/e16020953.

[19] Natal J, Ávila I, Tsukahara VB, Pinheiro M, Maciel CD. Entropy: from thermodynamics to information processing. Entropy 2021;23:1340. 10.3390/e23101340.

[20] Donoghue T, Hammonds R, Lybrand E, Washcke L, Gao R, Voytek B. Evaluating and comparing measures of aperiodic neural activity 2024:2024.09.15.613114. 10.1101/2024.09.15.613114.

[21] Lau ZJ, Pham T, Chen SHA, Makowski D. Brain entropy, fractal dimensions and predictability: a review of complexity measures for eeg in healthy and neuropsychiatric populations. Eur J Neurosci 2022;56:5047–69. 10.1111/ejn.15800.

[22] Waschke L, Wöstmann M, Obleser J. States and traits of neural irregularity in the age-varying human brain. Sci Rep 2017;7:17381. 10.1038/s41598-017-17766-4.

[23] Medel V, Irani M, Crossley N, Ossandón T, Boncompte G. Complexity and 1/f slope jointly reflect brain states. Sci Rep 2023;13:21700. 10.1038/s41598-023-47316-0.

[24] Stam CJ. Nonlinear dynamical analysis of EEG and MEG: Review of an emerging field. Clin Neurophysiol 2005;116:2266–301. 10.1016/j.clinph.2005.06.011.

[25] Frohlich J, Ruch S, Trunk BH, Keute M, Mediano PAM, Gharabaghi A. Brain signal complexity and aperiodicity predict human corticospinal excitability 2024:2024.02.09.579457. 10.1101/2024.02.09.579457.

[26] Roberts SJ, Penny W, Rezek I. Temporal and spatial complexity measures for electroencephalogram based brain-computer interfacing. Med Biol Eng Comput 1999;37:93–8. 10.1007/BF02513272.

[27] Krishnan PT, Joseph Raj AN, Balasubramanian P, Chen Y. Schizophrenia detection using multivariate empirical mode decomposition and entropy measures from multichannel EEG signal. Biocybern Biomed Eng 2020;40:1124–39. 10.1016/j.bbe.2020.05.008.

[28] Ashokkumar SR, Anupallavi S, Premkumar M, Jeevanantham V. Implementation of deep neural networks for classifying electroencephalogram signal using fractional S-transform for epileptic seizure detection. Int J Imaging Syst Technol 2021;31:895–908. 10.1002/ima.22565.

[29] Lal U, Chikkankod AV. Leveraging SVD entropy and explainable machine learning for Alzheimer’s and frontotemporal dementia detection using eeg 2023. 10.36227/techrxiv.23992554.v1.

[30] Belyaev M, Murugappan M, Velichko A, Korzun D. Entropy-based machine learning model for fast diagnosis and monitoring of parkinson’s disease. Sensors 2023;23:8609. 10.3390/s23208609.

[31] Muncaster ARG, Sleigh JW, Williams M. Changes in consciousness, conceptual memory, and quantitative electroencephalographical measures during recovery from sevoflurane-and remifentanil-based anesthesia. Anesth Analg 2003;96:720. 10.1213/01.ANE.0000040143.95962.36.

[32] Lal U, Mathavu Vasanthsena S, Hoblidar A. Temporal feature extraction and machine learning for classification of sleep stages using telemetry polysomnography. Brain Sci 2023;13:1201. 10.3390/brainsci13081201.

[33] Veyrié A, Noreña A, Sarrazin J-C, Pezard L. Information-theoretic approaches in EEG correlates of auditory perceptual awareness under informational masking. Biology 2023;12:967. 10.3390/biology12070967.

[34] Kiers L, Cros D, Chiappa KH, Fang J. Variability of motor potentials evoked by transcranial magnetic stimulation. Electroencephalogr Clin Neurophysiol Potentials Sect 1993;89:415–23. 10.1016/0168-5597(93)90115-6.

[35] Rossi S, Antal A, Bestmann S, Bikson M, Brewer C, Brockmöller J, et al. Safety and recommendations for TMS use in healthy subjects and patient populations, with updates on training, ethical and regulatory issues: Expert Guidelines. Clin Neurophysiol 2021;132:269–306. 10.1016/j.clinph.2020.10.003.

[36] Mills KR, Boniface SJ, Schubert M. Magnetic brain stimulation with a double coil: the importance of coil orientation. Electroencephalogr Clin Neurophysiol Potentials Sect 1992;85:17–21. 10.1016/0168-5597(92)90096-T.

[37] Sorensen HV, Burrus CS, Jones DL. A new efficient algorithm for computing a few DFT points. 1988 IEEE Int. Symp. Circuits Syst. ISCAS, 1988, p. 1915–8 vol.2. 10.1109/ISCAS.1988.15312.

[38] Gramfort A, Luessi M, Larson E, Engemann DA, Strohmeier D, Brodbeck C, et al. MEG and EEG data analysis with MNE-Python. Front Neurosci 2013;7. 10.3389/fnins.2013.00267.

[39] Li A, Feitelberg J, Saini AP, Höchenberger R, Scheltienne M. MNE-ICALabel: Automatically annotating ICA components with ICLabel in Python. J Open Source Softw 2022;7:4484. 10.21105/joss.04484.

[40] Pion-Tonachini L, Kreutz-Delgado K, Makeig S. ICLabel: An automated electroencephalographic independent component classifier, dataset, and website. NeuroImage 2019;198:181–97. 10.1016/j.neuroimage.2019.05.026.

[41] Barascud N, Alcocer P, Roujansky P, Tresols JJT, Szul M, Pals M, et al. nbara/python-meegkit: v0.1.3 2022.

[42] Appelhoff S, Hurst AJ, Lawrence A, Li A, Mantilla Ramos YJ, O’Reilly C, et al. PyPREP: A Python implementation of the preprocessing pipeline (PREP) for EEG data. 2022. 10.5281/zenodo.6363576.

[43] Jas M, Engemann DA, Bekhti Y, Raimondo F, Gramfort A. Autoreject: Automated artifact rejection for MEG and EEG data. NeuroImage 2017;159:417–29. 10.1016/j.neuroimage.2017.06.030.

[44] Vallat R. AntroPy: entropy and complexity of time-series in Python 2022.

[45] Rogasch NC, Sullivan C, Thomson RH, Rose NS, Bailey NW, Fitzgerald PB, et al. Analysing concurrent transcranial magnetic stimulation and electroencephalographic data: A review and introduction to the open-source TESA software. NeuroImage 2017;147:934–51. 10.1016/j.neuroimage.2016.10.031.

[46] Bigdely-Shamlo N, Mullen T, Kothe C, Su K-M, Robbins KA. The PREP pipeline: standardized preprocessing for large-scale EEG analysis. Front Neuroinformatics 2015;9:16. 10.3389/fninf.2015.00016.

[47] de Cheveigné A, Arzounian D. Robust detrending, rereferencing, outlier detection, and inpainting for multichannel data. NeuroImage 2018;172:903–12. 10.1016/j.neuroimage.2018.01.035.

[48] R Core Team. R: A language and environment for statistical computing. 2021.

[49] Jolly E. Pymer4: connecting R and python for linear mixed modeling. J Open Source Softw 2018;3:862. 10.21105/joss.00862.

[50] Bates D, Mächler M, Bolker B, Walker S. Fitting linear mixed-effects models using lme4. J Stat Softw 2015;67:1–48. 10.18637/jss.v067.i01.

[51] Kuznetsova A, Brockhoff PB, Christensen RHB. lmerTest package: tests in linear mixed effects models. J Stat Softw 2017;82:1–26. 10.18637/jss.v082.i13.

[52] Benjamini Y, Yekutieli D. The control of the false discovery rate in multiple testing under dependency. Ann Stat 2001;29:1165–88. 10.1214/aos/1013699998.

[53] Darling WG, Wolf SL, Butler AJ. Variability of motor potentials evoked by transcranial magnetic stimulation depends on muscle activation. Exp Brain Res 2006;174:376–85. 10.1007/s00221-006-0468-9.

[54] Frohlich J, Chiang JN, Mediano PAM, Nespeca M, Saravanapandian V, Toker D, et al. Neural complexity is a common denominator of human consciousness across diverse regimes of cortical dynamics. Commun Biol 2022;5:1–17. 10.1038/s42003-022-04331-7.

[55] Hadra M, Omidvarnia A, Mesbah M. Temporal complexity of EEG encodes human alertness. Physiol Meas 2022;43:095002. 10.1088/1361-6579/ac8f80.

[56] Strauss M, Sitt JD, Naccache L, Raimondo F. Predicting the loss of responsiveness when falling asleep in humans. NeuroImage 2022;251:119003. 10.1016/j.neuroimage.2022.119003.

[57] Türker B, Musat EM, Chabani E, Fonteix-Galet A, Maranci J-B, Wattiez N, et al. Behavioral and brain responses to verbal stimuli reveal transient periods of cognitive integration of the external world during sleep. Nat Neurosci 2023:1–13. 10.1038/s41593-023-01449-7.

[58] Aljalal M, Aldosari SA, Molinas M, AlSharabi K, Alturki FA. Detection of Parkinson’s disease from EEG signals using discrete wavelet transform, different entropy measures, and machine learning techniques. Sci Rep 2022;12:22547. 10.1038/s41598-022-26644-7.

[59] Bosl WJ, Tager-Flusberg H, Nelson CA. EEG Analytics for Early Detection of Autism Spectrum Disorder: A data-driven approach. Sci Rep 2018;8:6828. 10.1038/s41598-018-24318-x.

[60] Čukić M, Stokić M, Simić S, Pokrajac D. The successful discrimination of depression from EEG could be attributed to proper feature extraction and not to a particular classification method. Cogn Neurodyn 2020;14:443–55. 10.1007/s11571-020-09581-x.

[61] Casali AG, Gosseries O, Rosanova M, Boly M, Sarasso S, Casali KR, et al. A theoretically based index of consciousness independent of sensory processing and behavior. Sci Transl Med 2013;5:198ra105-198ra105. 10.1126/scitranslmed.3006294.

[62] Darmani G, Nieminen JO, Bergmann TO, Ramezanpour H, Ziemann U. A degraded state of consciousness in healthy awake humans? Brain Stimul Basic Transl Clin Res Neuromodulation 2021;14:710–2. 10.1016/j.brs.2021.04.012.

[63] Ort A, Smallridge JW, Sarasso S, Casarotto S, von Rotz R, Casanova A, et al. TMS-EEG and resting-state EEG applied to altered states of consciousness: oscillations, complexity, and phenomenology. iScience 2023;26:106589. 10.1016/j.isci.2023.106589.

[64] Sarasso S, Boly M, Napolitani M, Gosseries O, Charland-Verville V, Casarotto S, et al. Consciousness and Complexity during Unresponsiveness Induced by Propofol, Xenon, and Ketamine. Curr Biol 2015;25:3099–105. 10.1016/j.cub.2015.10.014.

[65] Perez MA, Cohen LG. Interhemispheric inhibition between primary motor cortices: what have we learned? J Physiol 2009;587:725–6. 10.1113/jphysiol.2008.166926.

[66] Belardinelli P, König F, Liang C, Premoli I, Desideri D, Müller-Dahlhaus F, et al. TMS-EEG signatures of glutamatergic neurotransmission in human cortex. Sci Rep 2021;11:8159. 10.1038/s41598-021-87533-z.

[67] Premoli I, Rivolta D, Espenhahn S, Castellanos N, Belardinelli P, Ziemann U, et al. Characterization of GABAB-receptor mediated neurotransmission in the human cortex by paired-pulse TMS–EEG. NeuroImage 2014;103:152–62. 10.1016/j.neuroimage.2014.09.028.

[68] Krile L, Ensafi E, Cole J, Noor M, Protzner AB, McGirr A. A dose-response characterization of transcranial magnetic stimulation intensity and evoked potential amplitude in the dorsolateral prefrontal cortex. Sci Rep 2023;13:18650. 10.1038/s41598-023-45730-y.

[69] Darmani G, Bergmann TO, Zipser C, Baur D, Müller-Dahlhaus F, Ziemann U. Effects of antiepileptic drugs on cortical excitability in humans: A TMS-EMG and TMS-EEG study. Hum Brain Mapp 2019;40:1276–89. 10.1002/hbm.24448.

[70] Khademi F, Royter V, Gharabaghi A. Distinct beta-band oscillatory circuits underlie corticospinal gain modulation. Cereb Cortex 2018;28:1502–15. 10.1093/cercor/bhy016.

[71] Schaworonkow N, Triesch J, Ziemann U, Zrenner C. EEG-triggered TMS reveals stronger brain state-dependent modulation of motor evoked potentials at weaker stimulation intensities. Brain Stimul Basic Transl Clin Res Neuromodulation 2019;12:110–8. 10.1016/j.brs.2018.09.009.

[72] Min J, Wang P, Hu J. Driver fatigue detection through multiple entropy fusion analysis in an EEG-based system. PLOS ONE 2017;12:e0188756. 10.1371/journal.pone.0188756.

[73] Xu R, Zhang C, He F, Zhao X, Qi H, Zhou P, et al. How physical activities affect mental fatigue based on eeg energy, connectivity, and complexity. Front Neurol 2018;9.

[74] Zhang T, Chen J, He E, Wang H. Sample-entropy-based method for real driving fatigue detection with multichannel electroencephalogram. Appl Sci 2021;11:10279. 10.3390/app112110279.

[75] Young JH, Arterberry ME, Martin JP. Contrasting electroencephalography-derived entropy and neural oscillations with highly skilled meditators. Front Hum Neurosci 2021;15.

[76] Karabanov AN, Madsen KH, Krohne LG, Siebner HR. Does pericentral mu-rhythm “power” corticomotor excitability? – A matter of EEG perspective. Brain Stimulat 2021;14:713–22. 10.1016/j.brs.2021.03.017.

[77] Khademi F, Royter V, Gharabaghi A. State-dependent brain stimulation: Power or phase? Brain Stimulat 2019;12:296–9. 10.1016/j.brs.2018.10.015.

[78] Hussain SJ, Claudino L, Bönstrup M, Norato G, Cruciani G, Thompson R, et al. Sensorimotor oscillatory phase–power interaction gates resting human corticospinal output. Cereb Cortex 2019;29:3766–77. 10.1093/cercor/bhy255.

[79] Liu P, Song D, Deng X, Shang Y, Ge Q, Wang Z, et al. The effects of intermittent theta burst stimulation (iTBS) on resting-state brain entropy (BEN). Neurotherapeutics 2025:e00556. 10.1016/j.neurot.2025.e00556.

[80] Huber R, Ghosh A. Large cognitive fluctuations surrounding sleep in daily living. iScience 2021;24:102159. 10.1016/j.isci.2021.102159.

[81] Valdez P. Circadian rhythms in attention. Yale J Biol Med 2019;92:81–92.

[82] Andrillon T, Burns A, Mackay T, Windt J, Tsuchiya N. Predicting lapses of attention with sleep-like slow waves. Nat Commun 2021;12:3657. 10.1038/s41467-021-23890-7.

[83] Noreika V, Kamke MR, Canales-Johnson A, Chennu S, Bekinschtein TA, Mattingley JB. Alertness fluctuations when performing a task modulate cortical evoked responses to transcranial magnetic stimulation. NeuroImage 2020;223:117305. 10.1016/j.neuroimage.2020.117305.

[84] Xu Y, Wokke M, Noreika V, Bareham C, Jagannathan S, Georgieva S, et al. Effects of alertness on perceptual detection and discrimination 2023:2023.03.21.533623. 10.1101/2023.03.21.533623.

[85] Nir Y, Andrillon T, Marmelshtein A, Suthana N, Cirelli C, Tononi G, et al. Selective neuronal lapses precede human cognitive lapses following sleep deprivation. Nat Med 2017;23:1474–80. 10.1038/nm.4433.

[86] Keerativittayayut R, Aoki R, Sarabi MT, Jimura K, Nakahara K. Large-scale network integration in the human brain tracks temporal fluctuations in memory encoding performance. eLife 2018;7:e32696. 10.7554/eLife.32696.

